# Transposable elements as evolutionary substrates of protein disorder in the human proteome

**DOI:** 10.64898/2026.06.12.731867

**Authors:** Juan Mac Donagh, Nicolas Vergesio, Andrea Aguilar, Rodrigo Nores, Antonio Lagares, Maria Silvina Fornasari, Gustavo Parisi

## Abstract

Intrinsically disordered regions (IDRs) are central contributors to protein function, evolution and human disease, yet the evolutionary routes that seed new disordered segments within pre-existing proteins are still poorly understood. Sequence insertions provide a powerful mechanism for disorder expansion, but the genomic donors of inserted IDR and its long-term conformational fate remain largely unknown. Transposable elements (TEs), abundant mobile genetic elements with distinctive compositional biases, represent compelling candidates for generating disorder within proteins. Here, we systematically mapped TE-derived segments across human proteins and isoforms, and we found that these insertions are strongly enriched in intrinsic disorder. The structural consequences of their insertion are shaped by TE class and family, reflecting the sequence biases of the elements from which they originate. Recent, Primate-specific insertions preferentially generate disordered segments, whereas older insertions more frequently occupy ordered structural contexts, revealing an age-dependent transition in the conformational state of TE-derived sequences. TE-containing isoforms are expressed at lower levels than TE-free isoforms, particularly when insertions are young and disorder-rich, suggesting that intrinsic disorder may constrain the cellular tolerance of newly exonized sequences. These findings identify TEs as a major evolutionary mechanism linking genome mobility to the emergence of new disordered conformational ensembles in the human proteome.

## Introduction

Intrinsic disorder confers proteins with a remarkable conformational plasticity, allowing the exploration of conformational landscapes that remain inaccessible to ordered proteins ^1–4^. Consistent with their higher evolutionary rates, intrinsically disordered regions (IDRs) act as permissive evolutionary substrates that readily accommodate insertions and point mutations ^5,6^. As a result, they serve as powerful vehicles for evolutionary innovation and functional diversification ^7^. Although IDRs display remarkable conformational flexibility, the term encompasses a wide spectrum of structural behaviors with distinct functional implications, ranging from highly expanded ensembles to more compact disordered conformations, as well as segments that fold upon interaction with their partners and others that remain disordered even in the bound state ^8–11^. Despite their well-characterized conformational properties and central roles in human biology and disease, the mechanisms governing IDR emergence and evolutionary trajectories remain poorly understood. Sequence insertions are widely recognized as a major driver of disorder expansion, yet little is known about where these inserted segments come from, what sequence and compositional properties they contribute, and how those properties determine the structural behavior of the newly formed region. Equally unclear is the long-term evolutionary fate of such insertions: whether they are retained as disordered elements or progressively reshaped into ordered structural domains under selective pressure.

Transposable elements (TEs) are a particularly compelling candidate source of such insertions: they are abundant, mobile, and have repeatedly contributed coding and regulatory material to host genomes throughout evolution. They populate eukaryotic genomes and propagate through repeated insertion events ^12–14^. TE insertions have been extensively investigated over the past two decades for their roles in genome evolution and stability ^15–18^, aging and human disease ^19–23^, the modulation of gene regulatory networks ^24,25^, and the generation of novel isoforms and protein neofunctionalization ^26–29^.

Here, we investigated the extent to which TE insertions have contributed to the emergence of intrinsically disordered regions in human proteins. We systematically identified and structurally mapped TE-derived segments across the human proteome to assess their contribution to the emergence of novel protein sequences and conformational landscapes. Focusing on major Primate-specific transposable element families, including SINEs (Short Interspersed Nuclear Elements), primarily Alu elements ^30^, Retroposon elements, mainly SVA (SINE-VNTR-Alu) ^31,32^, selected LINE (Long Interspersed Nuclear Elements) ^15^, and DNA transposon families, we examined how their insertions have shaped protein sequence and structure. Consistent with previous studies, TE insertions within genes are widespread and have been implicated in multiple cases of protein neofunctionalization through exonization ^26,27,33–38^, for a recent review, see ^39^. Building on this framework, we show that the conformational behavior of TE-derived protein segments is strongly influenced by the type of transposable element from which they originate. While the majority of TE-derived sequences give rise to IDRs, reflecting their compositionally biased sequence features, these regions are not evolutionarily static. Both TE-derived segments and their flanking regions, initially disordered upon insertion, can progressively transition toward more ordered structural states. Consistently, by leveraging estimated insertion ages across TE families, we find that evolutionarily recent TE insertions are strongly associated with intrinsic disorder, whereas older insertions more frequently overlap ordered regions.

TE insertions in CDS (coding sequences), therefore, emerge as an important mechanism for generating conformational novelty, capable of abruptly reshaping protein dynamics and ensemble behaviour in ways that differ fundamentally from the incremental effects of point mutations. Together, our findings identify transposable element insertions as a major and temporally traceable source of intrinsically disordered regions in human proteins, providing a mechanistic framework to study the origin, evolution, and functional diversification of protein disorder.

### Prediction of TE insertions in human CDS

To identify TE insertions within human CDSs we employed two complementary approaches. First, we used the Dfam database ^40^ with nhmmer ^41^ to screen for protein-coding human spliced transcripts derived from the Ensembl database (GRCh38.p14). Secondly, we downloaded precomputed RepeatMasker annotations from Dfam containing genome-wide mapped TE insertions (see Methods and Extended Data Fig. 1a). Using a reference set of 20,659 human protein-coding genes from UniProt we found that 92.20% of human genes harbor at least one sequence stretch with sequence similarity to TEs, but most of the TE sequences occur within introns (99.19%, Fig. 1a,b). From the remaining exonic regions, only 10.22% belongs to CDS (Fig. 1c, Supplementary Table 1–2). After applying different filters (see Methods) we identified 6,432 transcripts containing 7,039 TE insertions within coding regions across 3,511 genes, including both canonical proteins and isoforms according to UniProt (Supplementary Tables 2 and 3, Fig. 1d,e). The total number of insertions corresponds to 918 Dfam entries, where 24.07% (221) corresponds to Class II (DNA transposons), while 75.16% (690) belong to Class I (retrotransposons), with the remaining 0.77% corresponding to unclassified classes of TEs. Among the most abundant groups, elements from the SINE subfamily, followed by LTR (Long-Terminal Repeats) and LINE, were the most frequent insertions (Fig. 1f). The mean number of different insertions per transcript was 1.1, with most TE-derived segments encoded within a single exon and average length of 73 amino acids (Extended Data Fig. 1b–1d). TEs insertion distribution likely reflects the exceptionally high copy number of Alu elements (*>*1 million copies ^42,43^) in the human genome, which makes them the most frequent transposable element family despite their smaller contribution to the total genomic sequence. Also, as previously shown ^43,44^, we found that Alu elements display an overall bias toward the negative strand, while the other TE classes show a significant preference for positive strand insertions (Extended Data Fig. 1e, Supplementary Table 3). Finally, we found that 27.0% of the total TE insertions in CDS correspond to Primate-specific transposable elements ^45–48^ (Supplementary Table 4).

**Figure 1.**
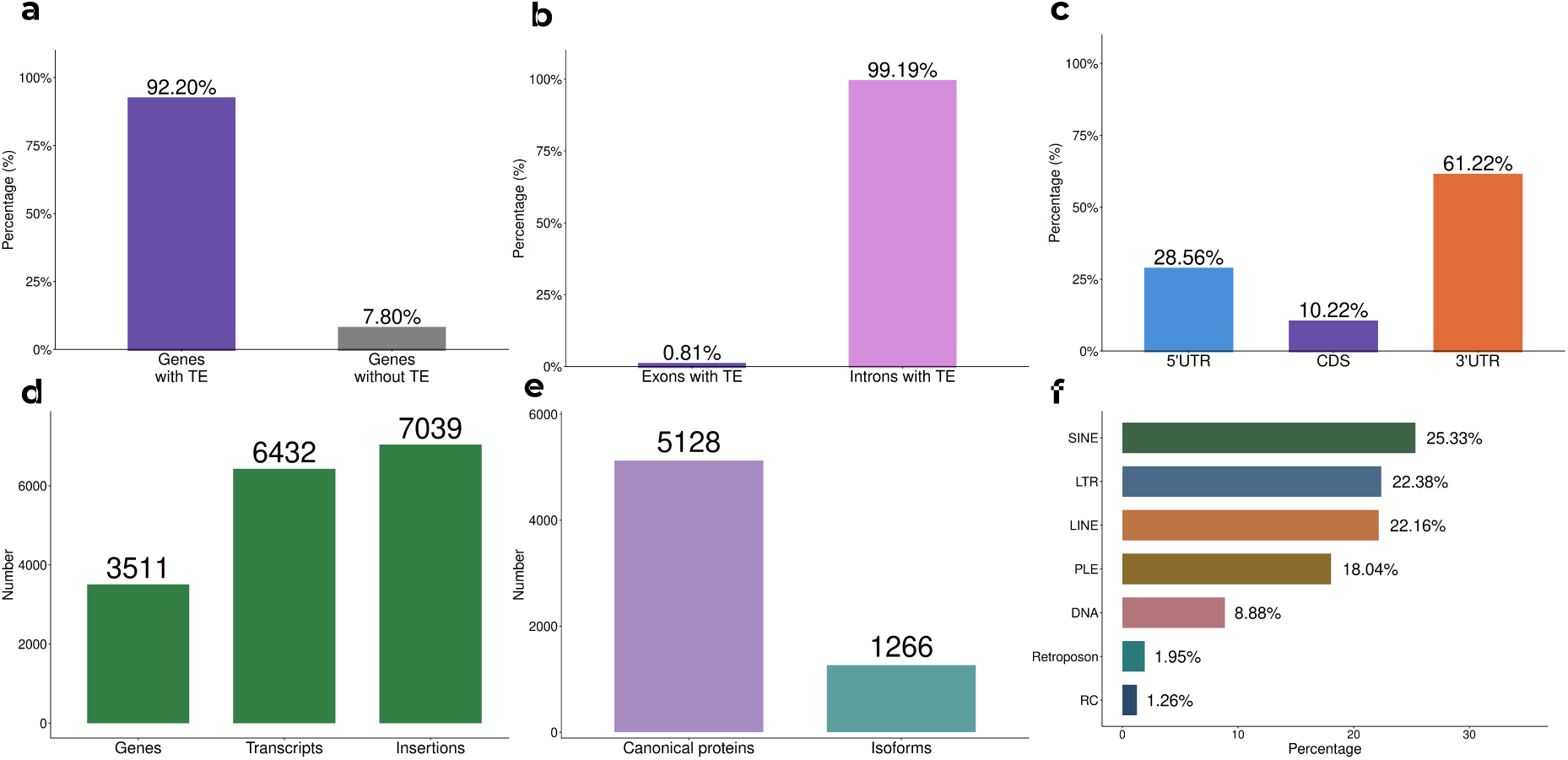
Transposable element insertions in the human proteome: **a.** Percentage of human genes containing at least one TE insertion. **b.** Distribution of TE insertions across exonic and intronic regions. **c.** Distribution of TE insertions across transcript regions. **d.** Total number of unique genes, transcripts and insertions identified. **e.** Number of canonical proteins and isoforms harboring TE insertions. **f.** Distribution of TE insertions by repeat type.

Beyond confirming the pervasive presence of TEs in the protein-coding fraction of the human genome in agreement with previous works ^27^, the breadth of our mapping enables the systematic structural and evolutionary analyses presented below, to deepen our understanding on the origin of disorder in the human proteome.

### TE insertions in coding regions are enriched in intrinsic disorder

All Ensembl transcripts used in the detection of TE insertions were mapped to UniProt to retrieve coding sequences and corresponding AlphaFold structural models (n = 69,449 protein sequences; 37,743 AlphaFold2 structures ^49^). To choose which predictors we used for IDRs detection, we benchmarked five different approaches: AIUPred ^50^, AlphaFold-RSA ^51^, AlphaFold pLDDT (1 − pLDDT), SPOT-Disorder ^52^, and PUNCH2-Light ^53^. Method performance was tested against a curated reference dataset using ROC analysis following CAID evaluation standards ^54^. Based on performance evaluation (see Methods and Fig. 2a, Supplementary Table 5) the three best-performing predictors (AlphaFold 1 − pLDDT, AUC = 0.811, AlphaFold-RSA, AUC = 0.801, and PUNCH2-Light, AUC = 0.806) were used to estimate disorder content across all translated transcripts, stratifying the dataset into proteins with and without TE insertions. As alternative isoforms have been proposed to exhibit higher disorder content than canonical proteins ^55^, we further partitioned the remaining dataset into canonical proteins, and UniProt annotated isoforms (Supplementary Table 6). Proteins harbouring TE-derived segments were significantly more disordered, on average, than both canonical proteins and annotated isoforms, reflecting a consistent enrichment in intrinsically disordered regions (Fig. 2b, Mann–Whitney U, r = 0.20 vs isoforms and r = 0.25 vs canonical proteins, both p *<* 0.0001; exact p values size effects, mean and medians are available in Supplementary Table 13 and Supplementary Table 14). This enrichment was robust across the three top-performing predictors (Extended Data Fig. 2a–2b). TE-containing proteins displayed the highest proportion of IDPs and highly disordered proteins when each group was compared to itself, among all three groups (Fig. 2c), representing 18.4% of the total amount of IDP in the human proteome (Extended Data Fig. 2c). Position-specific disorder analysis showed that, although both TE-derived and non-TE regions include ordered and disordered residues (Fig. 2d), TE derived regions are overall significantly more disordered than their non-TE counterparts, although the position-level effect across all proteins is small (Fig. 2d; r = 0.016, p *<* 0.0001, Extended Data Fig. 2d–2e, Supplementary Table 7).

**Figure 2.**
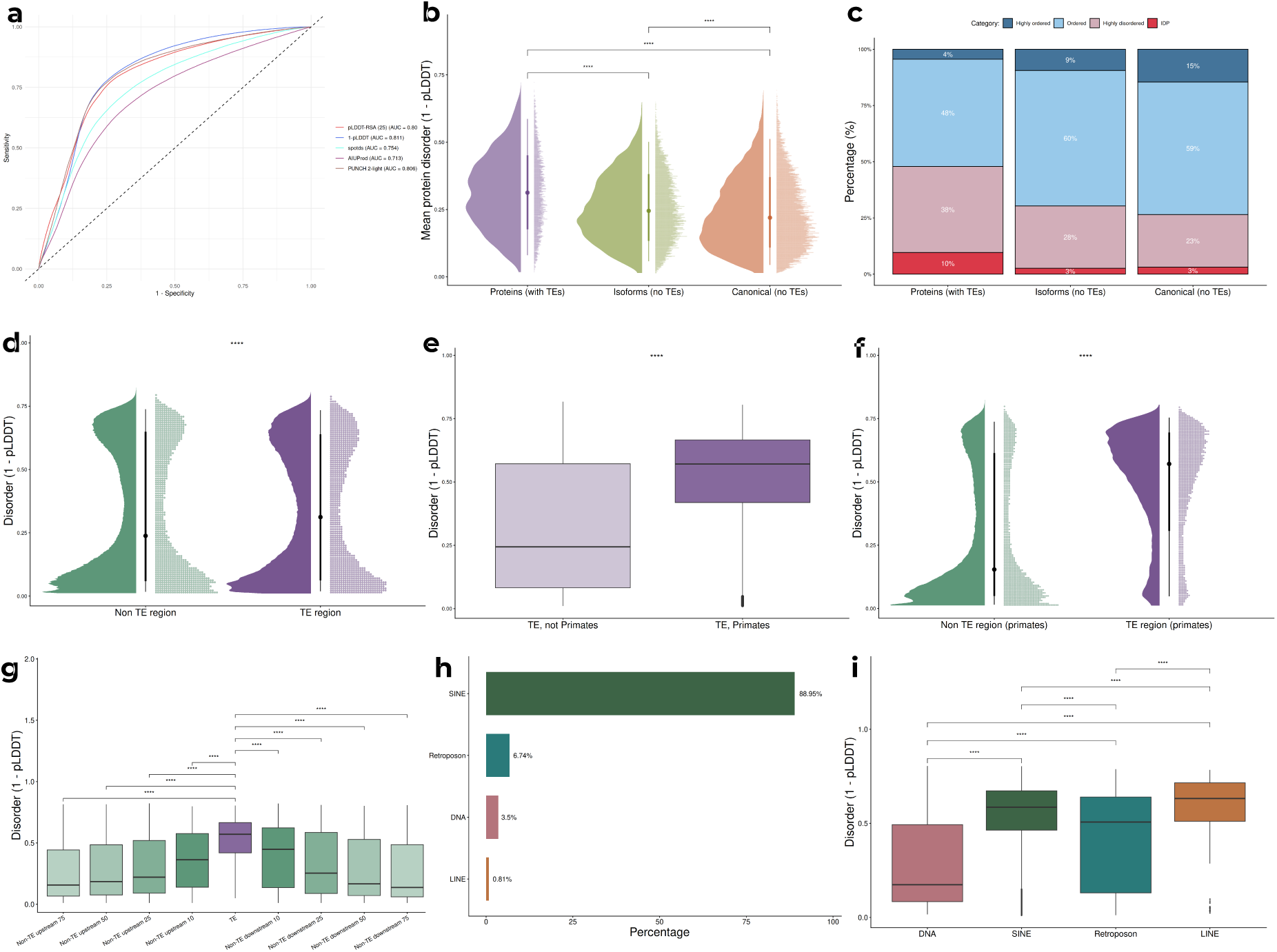
Transposable element insertions in coding sequences are enriched in intrinsic disorder: **a.** ROC curves for five disorder predictors benchmarked against a curated reference dataset (n = 1,913) following CAID evaluation standards. AUC values are shown in the legend. **b.** Distribution of mean disorder (1 − pLDDT) across three protein groups: proteins with TE insertions (n = 5,068), and isoforms or canonical proteins without TEs (n = 11,912 and n = 17,239). Proteins harbouring TE-derived segments are significantly more disordered than both control groups (Mann–Whitney U test, p *<* 0.0001). **c.** Proportions of disorder categories per group, classified by mean 1 − pLDDT values: highly ordered (*<* 0.10), ordered (0.10–0.321), highly disordered (0.321–0.5), and IDP (≥ 0.5). **d.** Position-specific disorder values comparing TE-derived and non-TE regions within proteins harbouring TE insertions (Mann–Whitney U test, p *<* 0.0001). **e.** Mean disorder of Primate versus Pre-Primate TE insertions (Mann–Whitney U test, p *<* 0.0001). **f.** Disorder distribution comparing TE-derived and flanking non-TE regions within Primate insertion proteins (Mann–Whitney U test, p *<* 0.0001). **g.** Disorder values of TE-derived regions compared to flanking non-TE segments at increasing distances (±10 to ±75 amino acids). **h.** Composition of Primate TE insertions by repeat type. **i.** Disorder distributions across TE classes for Primate TE insertions (DNA: 39 AlphaFold2 models with 3927 positions, Retroposon: 78 AlphaFold2 models and 2532 positions, SINE: 997 AlphaFold2 models and 37283 positions, LINE: 9 AlphaFold2 models and 288 positions). Boxplots show median and interquartile range; significance assessed by Mann–Whitney U test with Benjamini–Hochberg correction. ns, not significant; * p *<* 0.05; ** p *<* 0.01; *** p *<* 0.001; **** p *<* 0.0001.

TE insertions were then partitioned into Primate-specific (for brevity, Primate, TE lineages originated in Primates representing recent insertions, composing 27.0% of the initial dataset) and Pre-Primate-specific (for brevity, Pre-Primate TE lineages originated in other mammals representing older insertions, 38.5%, see Methods). We found that disorder content differs significantly between the groups: Primate insertions are more disordered than insertions predating the Pre-Primate lineage (Fig. 2e; r = 0.25, p *<* 0.0001). Furthermore, Primate TE insertions are markedly more disordered than their surrounding protein regions, recapitulating the overall trend observed across the full TE dataset but with an even stronger bias toward disorder (Fig. 2f; r = 0.26, p *<* 0.0001).

Primate insertions are overwhelmingly dominated by SINEs, which account for 88.78% of all events (Fig. 2h), primarily driven by AluS elements (57.7% of total insertions; Extended Data Fig. 3a). Comparative analysis across TE classes with all three predictors (Fig. 2i, Extended Data Fig. 3b–3c) revealed distinct structural behaviours: SINEs are preferentially associated with disorder-rich environments (mean 1 − pLDDT = 0.5431) and LINEs also showed elevated disorder values, though they have the smallest number of CDS-inserted TEs in Primates (mean 1 − pLDDT = 0.5846), whereas DNA transposons (mean 1 − pLDDT = 0.2818) display a more ordered pattern of insertion. In contrast, Retrotransposon elements exhibit a bimodal distribution of disorder (mean 1 − pLDDT = 0.4203), reflecting subtype-specific differences, even after redundancy was taken into account (see Methods): SVA E and SVA A are associated with lower 1 − pLDDT disorder values (0.3452 and 0.4081, respectively), whereas SVA C and SVA D show higher disorder values (0.6159 and 0.5747, respectively, Extended Data Fig. 3d). From a length perspective, DNA Primate display longer lengths (average of 101 amino acids) likely due to their enzymatic origin, while the rest of TE displayed lower and similar values (SINE: 37, Retroposon: 32 and LINE: 32 amino acids in average).

**Figure 3.**
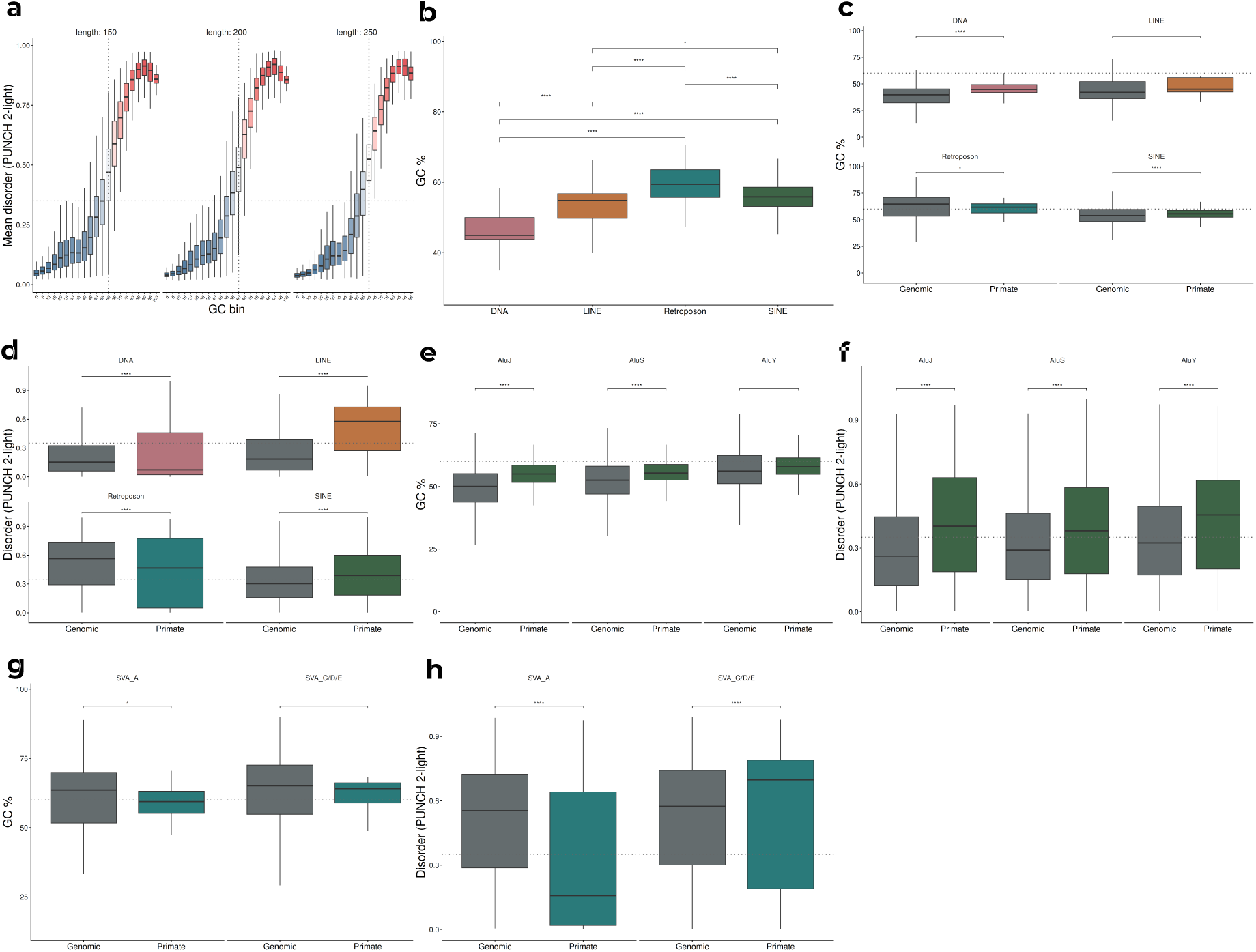
GC content and compositional properties drive disorder in transposable element-derived sequences: **a.** Mean disorder (PUNCH 2-light) of randomly generated sequences at varying GC content bins, stratified by sequence length (150, 200, and 250 amino acids). Disorder increases markedly above ∼55–60% GC content. Blue and red shading indicate ordered and disordered regimes respectively. **b.** GC content distribution of Primate TE-derived fragments per TE class (Mann–Whitney U test, p ≤ 0.001 for all comparisons). **c.** GC content comparison between genomic and Primate CDS-inserted fragments per TE class. **d.** Disorder comparison between genomic and Primate CDS-inserted fragments per TE class (Mann–Whitney U test, p *<* 0.0001). **e.** GC content comparison between genomic and Primate CDS-inserted Alu subfamilies (AluJ, AluS, AluY). **f.** Disorder comparison between genomic and Primate CDS-inserted Alu subfamilies. **g.** GC content comparison between genomic and Primate CDS-inserted SVA subfamilies (SVA_A and SVA_C/D/E). **h.** Disorder comparison between genomic and Primate CDS-inserted SVA subfamilies. Boxplots show median and interquartile range; significance assessed by Mann–Whitney U test with Benjamini–Hochberg correction. ns, not significant; * p *<* 0.05; ** p *<* 0.01; *** p *<* 0.001; **** p *<* 0.0001.

Given the hypothesis that exonization and insertion events can influence adjacent regions by promoting additional conformational changes, we next examined the structural properties of sequences flanking TE insertions. To further explore this, we analysed how the rest of the protein behaves in regard to the disorder value of the TE insertion, and we still found that TEs remain as the most disordered part of the protein after the insertion, but instead of examining the whole protein, we focused on the flanking regions of varying lengths from the TE. We defined segments of 10, 15, 25 and 75 amino acids from both the amino and carboxyl terminal boundaries of the inserted segments. Flanking regions followed the same general trend observed for inserted segments: TE insertion and the more closed region tended to be more disordered than those further away from the TE (Fig. 2g), effect sizes decreased monotonically with distance from the insertion (TE vs flank: r = 0.15 at ±10 aa rising to r = 0.46 at ±75 aa), consistent with a localized disordering effect that attenuates with distance. These findings suggest that TE insertions promote a localized disordering effect in the immediate structural neighbourhood of the insertion site.

Together, these findings suggest that TE-driven disorder is shaped by both element identity and evolutionary age. In some families, evolutionary time as derived from Primate and pre-Primates comparisons, appears to modulate the persistence of disorder, enabling a gradual transition from initially disordered segments toward more ordered structural states.

### Compositional determinants of TE-derived disorder

To investigate the basis of the association between TE-derived segments and protein disorder, we examined the GC content in CDS-inserted fragments. GC content in CDS has been previously linked to enrichment in disorder-promoting amino acids ^56,57^ and with the structural foldability of de novo genes ^58^. Following the protocol of Basile et al. ^57^, we generated random sequences spanning a range of GC contents. Using Punch-light, we observed that disorder becomes prevalent above ∼55–60% GC (Fig. 3a), consistent with previous findings. Figure 3b shows that GC of main TE families are compatible with their average composition. Alu elements are GC-rich, reflecting their origin as SINE retrotransposons derived from 7SL RNA ^17,59^ (Supplementary Table 8). Retroposon enrichment in GC similarly reflects their compositional architecture: they contain both an Alu-like region and a GC-rich variable number tandem repeat (VNTR) domain, features that favour the emergence of low-complexity, compositionally biased peptides associated with intrinsic disorder ^17,60^. In contrast, LINE- and DNA-transposon-derived fragments display lower GC content (Supplementary Table 9). These trends are consistent with the evolutionary constraints imposed by the protein-coding ORFs required for retrotransposition in LINEs ^61^ and by the transposases encoded by autonomous DNA TEs ^45^ which structural constraints on these ancestral coding sequences may persist even in highly degenerate copies. As derived from Fig. 2i, disorder is consistent with GC content for SINE, Retroposons and DNA transposons. LINE-derived CDS insertions deviate from this trend, showing more disorder than their GC content would predict. This anomaly is likely explained by their low occurrence in our dataset (n = 11 sequences).

Finally, we contrasted the disorder content of Primate TEs already inserted in CDS with the disorder that TE elements coming from the genome-wide pool of dispersed TEs would potentially encode if exonized (called here “Genomic TEs”), which vastly outnumbers the CDS-inserted set. To this end, we randomly selected sequences from the corresponding Dfam seed alignments, which were translated in all six reading frames (see Methods). We found that their GC contents are in global agreement with the fate of the inserted CDS fragments: DNA and LINE insertions would generate ordered fragments (38.69%, and 43.64%, respectively) while Retroposons and SINE disordered ones (62.76% and 52.92% respectively, Fig. 3c–3d).

At the TE subfamily level, the relationship between GC content and disorder is largely retained, but appears to be modulated by subfamily-specific features. Focusing on the most populated families, SINE (n = 1146) and Retroposon (n = 87) we found that SINE subfamilies, including AluJ, AluS, and AluY, followed the expected relationship between GC content and disorder (Fig. 3e–3f). Grouping Retroposons into older (SVA_A) and younger subfamilies (SVA_C/D/E) ^62^ to gain statistical power, revealed that both groups have GC contents high enough to favour disorder. However, older SVA members tended to be less disordered than younger members, mirroring the age-dependent disordering effect observed above (Fig. 3g–3h).

The tendency of younger Alu and SVA CDS insertions to harbor higher disorder may be partly explained by their GC-rich composition and by their high density of CpG dinucleotides, which are susceptible to methylation-dependent deamination. Over evolutionary time, this process drives CpG→TpG transitions, progressively eroding GC content as insertions age ^63^. Since GC-low codons are generally associated with structured, ordered protein regions, one might predict that older insertions generate increasingly ordered protein contexts ^57^.

Overall, TE-derived disorder appears to be compositionally encoded and evolutionarily modulated. GC-rich TE insertions, particularly Alu and SVA-derived fragments, provide a favourable substrate for the emergence of intrinsic disorder, whereas subsequent sequence erosion may gradually reduce this disordering potential over evolutionary time. Nevertheless, GC content alone does not fully explain TE-derived disorder. While this compositional feature is predictive in some TE families, in others disorder may also depend on protein-level constraints, including how the inserted segment is accommodated within the conformational and functional ensemble of the host protein.

### Disorder content of TE-derived segments transits to order with increasing insertion ages

To further explore the evolutionary behaviour of TE regions in CDS, we estimated insertion times for each Primate TE family (see Supplementary Table 10). The relative ages of each TE insertion were estimated using Kimura 2-parameter (K2P) distances ^64^ adjusted for CpG sites, calculated from RepeatMasker alignments using corresponding consensus from Dfam ^40^ for each Primate TE family. Sequence divergence from consensus serves as a proxy for insertion age, with lower values indicating more recent activity ^65^. Distances for TE regions in CDS for each family were compared to the corresponding distances of all TEs scattered along the human genome not involved in CDS insertions. At a global scale, the distribution of Kimura distances for all TEs within CDS overlaps to the genomic background (Fig. 4a). However, family-specific analyses revealed distinct patterns. While SINE and LINE inserted fragments show divergence profiles that mainly overlap with their genomic counterparts, DNA transposons are shifted toward lower divergence values compared to the rest of the genome. Notably, Retroposons display a bimodal distribution in the genome, but CDS insertions are restricted to a single peak of low divergence (Fig. 4b). This suggests that only relatively recent SVA insertions are maintained within coding regions, while older ones might have been purged or are less frequent. Further analysis of Alu subfamilies (AluJ, AluS, and AluY) confirmed that their divergence in CDS is in general consistent with their established chronological ages (Extended Data Figure 4a–b).

**Figure 4.**
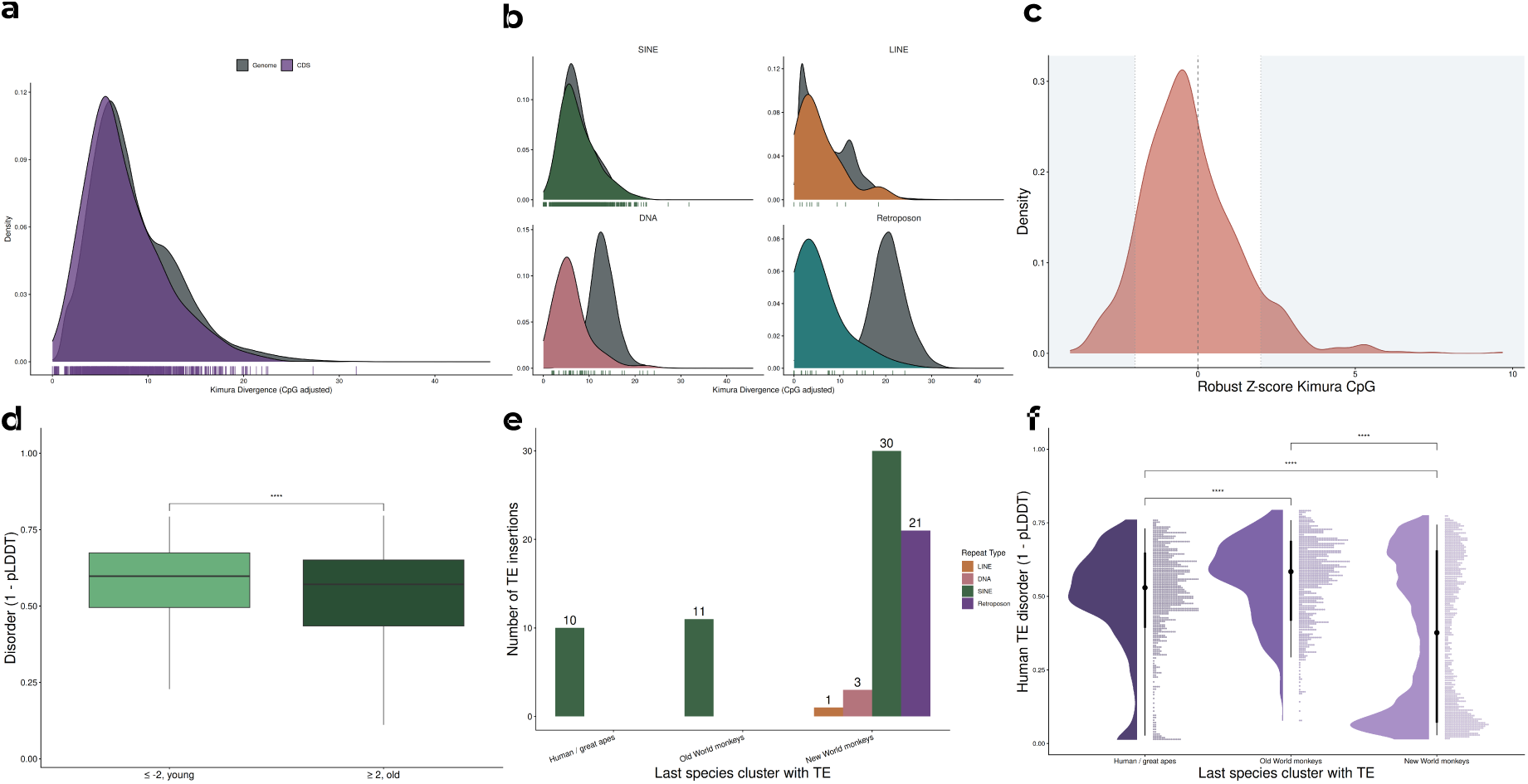
Evolutionary age of TE insertions correlates with disorder content in human proteins: **a.** Kimura CpG distribution for all inserted (CDS) transposons and all non-coding (Genomic) transposons. **b.** Kimura CpG distribution per specific TE type. **c.** Distribution of robust Z-scores of Kimura CpG divergence for all TE CDS insertions relative to the genome-wide distribution of the corresponding TE family. **d.** Disorder (1 − pLDDT) of Alu insertions with Z-score ≤ −2 (younger, n = 73 proteins) versus Z-score ≥ +2 (older, n = 81 proteins) (Mann–Whitney U test, p *<* 0.0001). **e.** Number of Primate TE insertions per repeat type, classified by the last Primate clade in which an orthologous insertion was detected (Human/great apes, Old World monkeys, New World monkeys). **f.** Disorder (1 − pLDDT) of human TE-derived segments classified by depth of orthologous conservation across Primate (pairwise Mann–Whitney U test, p *<* 0.0001). Boxplots show median and interquartile range; significance assessed by Mann–Whitney U test with Benjamini–Hochberg correction. ns, not significant; * p *<* 0.05; ** p *<* 0.01; *** p *<* 0.001; **** p *<* 0.0001.

Additionally, we identified CDS insertions whose Kimura distances deviated significantly from the corresponding genome-wide distribution of TE insertions of the same family, computing a Z-score for each CDS insertion relative to that genomic reference (Fig. 4c, Supplementary Table 11). Positive z-scores correspond to insertions older than the genomic average for the family, whereas negative z-scores correspond to younger insertions. In the well-sampled Alu set (n = 1,050 CDS insertions across AluS (57.70% of the Primate dataset), AluJ (26.45%) and AluY (4.63%)), we recovered 154 Alu elements that had either a Z-score ≤ −2 (younger, 73 proteins) or Z-score ≥ +2 (older, 81 proteins). Fig. 4d shows the distribution of 1 − pLDDT derived disorder for these two groups: younger Alu insertions exhibit significantly higher disorder than older ones (Fig. 4d; r = 0.12, p *<* 0.0001). Both AluJ and AluS showed the same tendency of younger insertions being more disordered than older ones (Mann–Whitney U p *<* 0.01), and AluY showed no significant difference (Extended Data Figure 5a).

Finally, we asked whether the disorder content of TE-derived segments is associated with their evolutionary age through orthology conservation across Primates. Because a human TE insertion present in distant Primate orthologs must have arisen before their last common ancestor, depth of orthologous conservation provides a phylogenetic proxy for insertion age that is independent of the sequence-based Kimura-distance approach used above. We compiled 604 orthologous sequences corresponding to 82 human proteins carrying Primate TE insertions (Fig. 4e), drawn from eleven Primate species spanning great apes (GA: *Homo sapiens*, *Pan troglodytes*, *Gorilla gorilla*, *Pongo abelii*), Old World monkeys (OWM: *Papio anubis*, *Macaca mulatta*, *Macaca fascicularis*, *Chlorocebus sabaeus*) and New World monkeys (NWM: *Callithrix jacchus*, *Saimiri boliviensis*, *Aotus nancymaae*). Aligned sequences were classified according to the most distant Primate clade in which an orthologous insertion was detectable (GA only, up to ∼18 MYA, to OWM, ∼30 MYA, or up to NWM, ∼40 MYA ^66^), and disorder in the human TE segment was compared across the three classes. Segments conserved only in GA and in OWM (i.e., the youngest insertions) showed significantly higher disorder than segments conserved into NWM (Fig. 4f; pairwise r = 0.16–0.32, all p *<* 0.0001; Supplementary Table 12, mean 1 − pLDDT: GA = 0.5247, OWM = 0.5747, NWM = 0.3364). Complementary to the Kimura-distance analysis presented above, this orthology-based result provides an independent line of evidence that older TE-derived insertions in human protein-coding sequences are systematically less disordered than younger ones.

Together, these findings demonstrate, using complementary methods to assess both disorder content and insertion timing, that TE-derived sequences contribute to the emergence of intrinsically disordered regions in the human proteome. Primate insertions are preferentially associated with disorder-promoting contexts, and evolutionary time modulates their structural fate: recent SINE insertions tend to retain high disorder content, while older insertions show a progressive shift toward more ordered conformations, consistent with gradual structural selection following exonization.

### TE-derived disorder in experimentally characterized proteins

To gain further support to our disorder predictions experimentally, we mapped all TE-containing human proteins against well-annotated curated databases of protein disorder such as DisProt ^67^ and MobiDB ^68^. As a result, 159 human proteins with TEs inserted in their CDS were found in the DisProt database, where 39 mapped to curated disordered regions. Among those with functional annotations we found Peroxiredoxin-4, an enzyme that catalyzes the reduction of hydrogen peroxide and organic hydroperoxides ^69^ and has an insertion of an LTR retrotransposon (ERV-like element) in positions 1 to 45 (Extended Data Figure 6a). According to the curated database Disprot, positions 1 to 75 are disordered. It has been described that Peroxiredoxin-4 in its oxidized state forms a homodecamer where the disordered N-terminal domain containing the TE insertion contributes to the stabilization of the complex. The mutant of Peroxiredoxin-4 containing a deletion of the first 46 N-terminal residues, dissociates into dimers after the reaction with H_2_O_2_, suggesting that the flexible region contributes to protein stabilization ^70^. Another example, the protein Nucleophosmin is a nucleolar protein in which the RNA binding activity plays essential roles in ribosome biogenesis ^71^. It has two main disordered regions, one composed by two acidic regions (120 to 188) and one basic (181 to 243). These regions have been characterized as disordered and are involved in the regulation of RNA binding ^72^. Precisely, the second acidic region has an Helitron insertion, a DNA transposon (positions 159 to 181) split into two consecutive exons (Extended Data Figure 6b). Helitrons have been shown to promote alternative splicing by introducing cryptic splice sites and by capturing host gene fragments, generating chimeric transcripts. A third example is the probable ATP-dependent RNA helicase, the human PRP28 (hPrp28, also known as DDX23) is phosphorylated by SRPK2, which is essential for the integration of the U4/U6-U5 tri-snRNP into the spliceosome ^73^. The amino terminal end of hPrp28 has a RS domain (Arginine-Serine rich domain, residues 1–138) containing key serine residues which upon phosphorylation promotes a conformational switch towards turn-like conformations ^74^. Amino acids from 70 to 140 from hPrp28 correspond to an insertion of a LINE element containing three out of ten of the phosphorylations sites (Extended Data Figure 6c).

Among proteins with TE insertions, 391 were annotated in MobiDB ^68^, which reports missing-residue regions in UniProt proteins, used here as a proxy for intrinsic disorder. Of these, 188 showed overlap between TE insertions and missing-residue regions (≥5% of the TE fragment). Rather than representing an upper limit of TE contribution to disorder, this proportion reflects the dual evolutionary fate of TE insertions: while many contribute to disordered regions, a substantial subset, particularly ancient insertions, has been incorporated into ordered protein regions as shown before. Actually, insertions of TE belonging to the same families originated ordered, disordered and also order-disorder transitions regions in different proteins with estimated different insertion times. Glyoxalase domain-containing protein 4, an enzyme recently associated with the nitration of alpha-synuclein ^75,76^ has an AluS insertion between positions 30 to 52. This region forms a flexible loop, which appears to transit from ordered to disordered conformation just above the active site of the enzyme (Extended Data Figure 6d) ^76^. In protein 5’-AMP-activated protein kinase subunit gamma-1 we found an AluS insertion in a completely ordered conformation in all examined crystallographic structures of the protein. Contrarily, there are cases within the AluJ family where TE insertions display ordered structures: the Parkin coregulated gene protein, with an AluJ insertion in positions 203 to 244, forms a large disordered loop which overlaps with 20 missing residues in PDBs. Similarly, an AluJ insertion (positions 269 to 289) in the tissue-type plasminogen activator forms a completely ordered PDB region in (Extended Data Figure 6e).

To further explore the conformational landscape of TE insertions in CDS, we mapped our dataset of proteins with the CoDNaS database ^77^, a database with experimentally determined conformational diversity of proteins derived from PDB. The protein mucosa-associated lymphoid tissue lymphoma translocation 1 (MALT1) is part of the CBM complex (CARD11–BCL10–MALT1), a key signaling hub that couples antigen receptor engagement to downstream activation of the transcription factor NF-κB ^78^. MALT1 has an insertion of a L1 element in positions 493 to 501. MALT1 is a protease whose active site is located in a pocket surrounded by 4 loops (L1 to L4) and a region called “elbow”. The L1 insertion mapped within part of the elbow and L3 carrying several amino acids that are crucial for substrate recognition (Extended Data Figure 6f). This region shows a large conformational change (16 Å) which is essential to stabilize the active conformation of the MALT1 ^79^.

### Expression profiles of TE-derived protein isoforms are evolutionary modulated

Transcriptional analysis of TE exonized proteins has been extensively studied in the last few years. Lin et al. ^27^ used a proteotranscriptomics approach to find evidence for the translational activities of 85 Alu exons in human proteins. More recently, Arribas et al. ^28^ showed that TE-exonizing isoforms are expressed at levels approximately tenfold lower than canonical transcripts, they are nonetheless efficiently translated, they are recurrent across individuals and functionally relevant despite their low expression. In line with these previous studies, we also found that transcripts harboring TE insertions showed, on average, lower expression levels than those without insertions (Supplementary Table 15, Fig. 5a). This global pattern, however, masks an underlying evolutionary signal: when insertions were stratified by age, older (pre-Primate) TEs exhibited significantly higher detectable expression than younger (Primate) ones (Fig. 5b). To minimize gene-specific regulatory effects on transcription, we focused on a set of 136 genes encoding at least one protein-coding isoform of each of the three classes: Primate TE-containing, Pre–Primate TE-containing, and TE-free isoforms (Supplementary Table 16). This within-gene comparison revealed a consistent hierarchy of expression levels, with Primate TE-containing isoforms exhibiting the lowest expression, followed by isoforms containing older TE insertions, and finally by isoforms lacking TE insertions, which showed the highest expression levels (Fig. 5c). Although no differences in disorder content was found considering the whole isoform lengths (Fig. 5d, Supplementary Table 16 and Supplementary Table 17), as described above, insertion fragments coming from Primate TE are significantly more disordered than Pre-Primate fragments (Fig. 5e, Supplementary Table 18).

**Figure 5.**
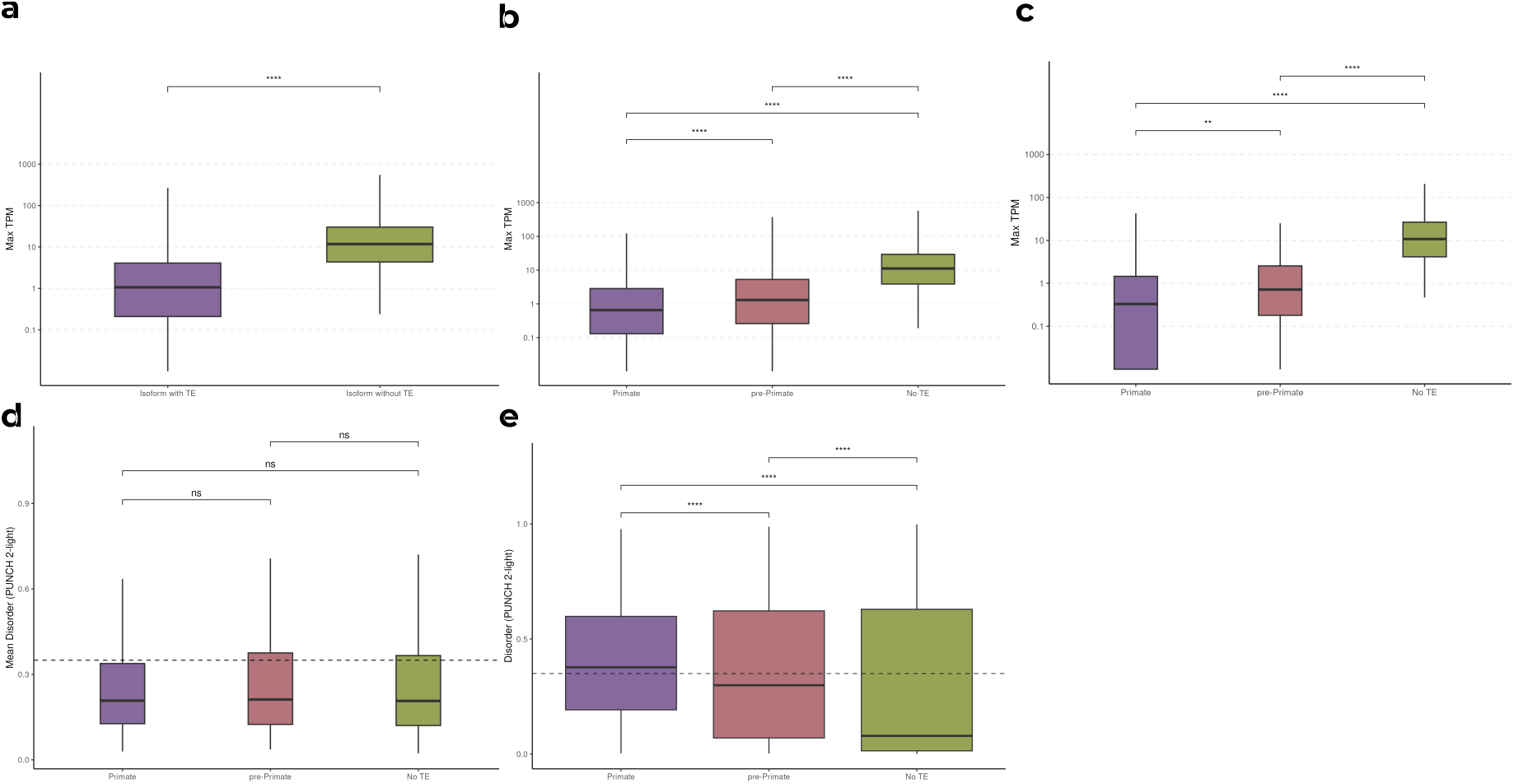
Expression and splicing properties of TE-containing isoforms. **a,** Maximum expression (max TPM across GTEx V10 tissues) of TE-containing versus TE-free isoforms (Wilcoxon rank-sum test). **b,** Expression stratified by TE evolutionary age across all isoforms. **c,** Within-gene matched comparison of expression levels across the three isoform classes in 136 genes encoding at least one representative of each class; significance assessed by Wilcoxon rank-sum test. **d,** Mean protein disorder for all 136 genes, grouped by Primate-specific, pre-Primate and TE-free transcripts (No TE). **e,** Predicted disorder per residue for transposon regions in Primate-Specific and pre-Primate genes, compared to all protein residues for transcripts without transposable elements (No TE). Unless otherwise noted, significance assessed by Mann–Whitney U test with Benjamini–Hochberg correction. ns, not significant; * p *<* 0.05; ** p *<* 0.01; *** p *<* 0.001; **** p *<* 0.0001.

In order to explain the observed expression profile differences between Primate and Pre-Primate containing isoforms, we explored 5^′^ and 3^′^ splice-site strengths for every internal exon in the three isoform classes using MaxEntScan ^80^ (Supplementary Table 19). We found that the strength of 5^′^ and 3^′^ were significatively higher for TE-free exons but no differences between Primate and Pre-Primate TE-containing exons were found (Extended Data Figure 7a–b). In the same line, we asked whether the differences might instead arise from the density of auxiliary cis-regulatory elements like exonic splicing enhancers and silencers (ESE/ESS) ^81,82^ using HEXplorer score ^83^, but again no differences between Primate and Pre–Primate were found (Extended Data Figure 7b–c, Supplementary Table 20).

Having excluded both gene-level regulatory differences, through our within-gene comparison, and local cis-regulatory features such as splice-site strength and enhancer density, we next asked whether the expression hierarchy could instead reflect differences in the structural properties of the encoded regions. Primate TE-derived segments are significantly more disordered than older TE-derived segments (Fig. 2e and 4e), raising the possibility that recently exonized, highly disordered peptide regions are subject to tighter expression control. Intrinsically disordered proteins are known to be tightly regulated from transcript synthesis to protein degradation ^84^, a constraint that arises in part because their conformational flexibility promotes promiscuous interactions making them particularly dosage-sensitive ^85,86^. Within this framework, the higher disorder content of Primate TE-derived segments may impose constraints on protein abundance that could partially explain the different expression profile compared with Pre-Primate TE insertions.

## Conclusions

Our results reframe transposable element insertions as one of the major mechanisms by which the human proteome acquires conformational novelty. By introducing compositionally biased and often disorder-prone segments into coding regions, TEs can abruptly generate new intrinsically disordered regions and locally perturb the structural behaviour of adjacent sequences. This mechanism differs from the gradual effects of point mutations and long evolutionary pathways because it introduces entire sequence modules whose conformational properties are already encoded in the composition of the TE class and family.

A central implication of our work is that protein disorder has a measurable evolutionary lifetime. We found that evolutionarily recently Primate insertions are enriched in disorder and tend to be incorporated into low-abundance isoforms, whereas older insertions more frequently occur in ordered contexts and show higher expression compatibility. Our results suggest that disordered TE-derived sequences may first be incorporated into the proteome as dosage-sensitive elements and subsequently undergo evolutionary filtering, regulatory accommodation or structural stabilization.

Together, these findings identify transposable elements as a major and temporally resolved source of protein disorder. More broadly, they suggest that mobile DNA elements have contributed not only to genome regulation and transcript diversification, but also to the origin of protein conformational ensembles, providing a mechanistic link between genome mobility and the emergence of new structural states in the human proteome.

## Acknowledgments

MSF, RN, AL, AA and GP are researchers and JMD and NV are PhD fellows from Consejo Nacional de Investigaciones Científicas y Técnicas (CONICET). This work was supported by Universidad Nacional de Quilmes (PUNQ 1004/11) and co-funded by the European Union under grant agreement no. 101182949 (HORIZON-MSCA-SE project IDPfun2); views and opinions expressed are, however, those of the author(s) only and do not necessarily reflect those of the European Union or the European Research Executive Agency. Neither the European Union nor the granting authority can be held responsible for them. This work used computational resources from UNC Supercómputo (CCAD) – Universidad Nacional de Córdoba (https://supercomputo.unc.edu.ar), which are part of SNCAD, República Argentina.

## Methods

### Computational environment

All analyses were performed in R version 4.5.3 (2026-03-11) running on a JupyterHub Linux. This work used computational resources from UNC Supercómputo (CCAD) – Universidad Nacional de Córdoba (https://supercomputo.unc.edu.ar), which are part of SNCAD, República Argentina. Data processing, statistical analyses and figure generation were carried out using ad hoc R scripts developed for this study, available at https://gitlab.com/sbgunq/publications/mac-donagh-te-2026. Effect sizes are reported as rank-biserial correlation r; given large sample sizes, we report r alongside p-values, with full per-comparison statistics in Supplementary Table 13 and Supplementary Table 14.

### Prediction of TE insertions in human CDS

All the coding sequences for protein-coding human transcripts were retrieved via BioMart, filtering by transcripts that are both encoded in a chromosomal region (taking out transcripts from scaffoldings). After this, the latest version of Dfam ^87^ (March 2025) was downloaded locally to a cluster, and we ran the HMMER suite ^88^ to detect annotated transposons in our transcripts. After filtering by e-value (*<* 0.005), a total of 8,690 transcripts were detected to hold one or more TEs. In addition to this, gene annotations (GTF files) were retrieved from Ensembl ^89^ (GRCh38.p14), and the genomic coordinates of protein-coding genes were used to retrieve transposable element (TE) hits from the Dfam API. Hits overlapping coding sequence (CDS) regions were retained using an e-value threshold of *<* 1 × 10^−10^ and requiring a minimum overlap of 30 nucleotides. This approach yielded an additional 1,877 transcripts containing TE-derived sequences. In total, we retrieved 10,522 unique transcripts that had 1,352 distinct transposable elements within their coding regions, which affected a total of 8,180 proteins (accounting for both canonical proteins and isoforms). To avoid redundancy, for transposons that had transposons mapped within the same region of nucleotides, we only choose the one with the lowest e-value and the largest length, so as not to map the same sequences twice, to finally obtain 6,432 unique transcripts, 7,039 TE insertions and 918 Dfam entries. (Distributions for e-value scores, final selected insertions per transcript, length of insertion and exon occupancy and the final numbers of genes, transcripts, unique transposons and types of protein are pictured in Figure 1).

### Prediction of disorder

To choose which predictors we used for IDRs detection, we benchmarked five different approaches: AIUPred ^90^, AlphaFold-RSA ^91^, AlphaFold pLDDT (1 − pLDDT), SPOT-Disorder ^92^, and PUNCH 2-Light ^93^. AlphaFold models were downloaded from https://alphafold.ebi.ac.uk/about. Method performance was benchmarked against a curated reference dataset (n = 1,913) derived from MobiDB ^94,95^ containing information about the presence of missing regions in PDB derived structures. This data was used to perform a ROC analysis following CAID evaluation standards ^96^.

### GC content and disorder

Following the protocol of Basile et al. ^97^, we generated random sequences (150, 200 and 250 amino acids in length) spanning a range of GC contents. These sequences were translated using the standard genetic code and protein disorder was predicted using PUNCH 2-light ^93^.

### Disordering content in genomic TEs

We randomly selected sequences from the corresponding Dfam seed alignment of main Primate TEs (SINE, Retroposons, DNA transposons and LINE1), which were translated in all six reading frames using the standard genetic code. When stop codons occurred, subsequences longer than 30 bp were retained. After this, disorder content was calculated using PUNCH 2-light ^93^.

### Partition of Primate specific, Pre-Primates and not Mammals TE insertions

We divided insertions according to a curated dataset containing TE origin with Dfam information from each entry (Supplementary Table 4). In order to reduce noise from our “background” dataset, we separated the Pre-Primate and not-Mammals entries and filtered out the latter: although these were retrieved mostly by our alignment method and carried annotated TE entries, they were annotated in species too distant from *Homo sapiens*. Entries specific to primates were hand-curated from the literature, available in Supplementary Table 3.

### TE insertion age estimation

#### Sequence alignment

To estimate the evolutionary divergence of CDS-inserted TE sequences relative to the genome-wide distribution of the same families, CDS-derived insertions and a matched set of genome-wide copies of the corresponding subfamilies were each aligned against the Dfam (March 2025 release) consensus sequence for each subfamily using RMBlastN (https://www.repeatmasker.org/rmblast/), with alignment parameters analogous to those applied by RepeatMasker (Smit, AFA, Hubley, R & Green, P. RepeatMasker Open-4.0. 2013–2015, http://www.repeatmasker.org). A relaxed E-value threshold (≤ 0.1) was applied to maximize sensitivity for low-copy families such as SVA_E. Per query, only the highest-scoring alignment was retained; full alignment parameters are available in the code repository https://gitlab.com/sbgunq/publications/mac-donagh-te-2026.

#### CpG-adjusted Kimura divergence

Evolutionary divergence was estimated with the two-parameter Kimura model (K2P) ^98^, incorporating an explicit correction for CpG hypermutability ^99^. Briefly, CG dinucleotides in the consensus sequence were treated as a single unit: coincident transitions were counted as one, single transitions were down-weighted to 0.1, and transversions were counted individually, with the effective alignment length adjusted accordingly. Full implementation is available in the code repository https://gitlab.com/sbgunq/publications/mac-donagh-te-2026.

#### Statistical analysis of divergence distributions

Divergence distributions were compared between CDS and genomic copies for each TE superfamily (SINE, LINE, Retroposon, and DNA). For Alu elements, distributions were further stratified by subfamily age class (AluJ, AluS, AluY). Normality of each distribution was assessed by Shapiro–Wilk test (subsampled to n = 5,000 when required), quantile–quantile plots, and skewness and excess kurtosis. To assess the divergence of each CDS insertion relative to the genome-wide background of its subfamily, robust Z-scores were computed as z = (x − M) / MAD, where M is the genomic median and MAD the median absolute deviation (scaled by the consistency constant 1.4826, as implemented in R’s mad() function) ^100^. CDS insertions were classified as relatively young (z ≤ −2) or relatively old (z ≥ +2) with respect to the genomic distribution of the corresponding subfamily.

#### Primates evolutionary conservation of TE-insertions

For the total set of Primate TE insertions, we looked for other Primate proteins carrying Primate TE insertions using the OMADB ^101^ database, drawn from eleven Primate species spanning great apes (GA) (*Homo sapiens*, *Pan troglodytes*, *Gorilla gorilla*, *Pongo abelii*), Old World monkeys (OWM) (*Papio anubis*, *Macaca mulatta*, *Macaca fascicularis*, *Chlorocebus sabaeus*) and New World monkeys (NWM) (*Callithrix jacchus*, *Saimiri boliviensis*, *Aotus nancymaae*). After each sequence had a set of homologues, we aligned each individual pair using MUSCLE ^102^, and looked for each corresponding region in the *Homo sapiens* TE. We compiled 604 orthologous sequences corresponding to 82 human proteins that had in at least one copy of the human TE. A TE insertion was considered present in a given species when the aligned region in that species contained no more than five percent of gap positions across its entire TE length. For each insertion, the most phylogenetically distant Primate retaining the TE-derived sequence was used to assign the insertion to one of three Primate clades. Each insertion was then annotated with its TE class (SINE, LINE, DNA, Retroposon) and the mean per-residue disorder (1 − pLDDT) of the TE-derived segment was compared across clades using Mann–Whitney U tests with Benjamini–Hochberg correction.

### Protein visualization

All protein structure visualizations were generated using UCSF ChimeraX version 1.10^103^. Structures were retrieved from the AlphaFold2 models and PDB entries associated with proteins from our initial dataset of TE-containing human transcripts. TE-derived segments and flanking regions were highlighted as described in the corresponding figure legends.

### Expression profiles and functional enrichment analysis

Gene-level expression was quantified from GTEx V10 RNA-seq data ^104^. For each gene harboring a TE-containing transcript, all isoforms of the same gene were extracted from the GTEx annotation, providing both TE-containing transcripts and their TE-free sister isoforms. Expression was summarized as the median TPM across all samples per tissue, and each transcript was then annotated with its maximum median TPM across the 30 GTEx tissues for downstream comparisons.

Transcripts were classified into four groups based on the evolutionary age of their TE insertions: Primate-specific (all TE hits Primate), Pre-Primate (all TE hits Pre-Primate), Not-mammal (all TE hits Not-mammals), mixed (transcripts harboring at least two different classes), and TE-free. For all subsequent analyses only protein-coding transcripts were considered and Not-mammals isoforms were excluded. For the global two-group comparison (TE-containing vs. TE-free), the most expressed isoform per group per gene was retained and differences in max_expression were assessed with a two-sided Wilcoxon rank-sum test. For the three-group comparison (Primate-TE, Pre-Primate-TE, no-TE), Not-mammal and mixed classes isoforms were excluded and Mann-Whitney U tests were performed with Benjamini–Hochberg (BH) correction. To control for gene-intrinsic regulatory differences, within-gene comparisons were restricted to the 136 genes encoding at least one isoform of each of the three classes; pairwise comparisons within this matched set were performed with BH correction.

Intrinsic disorder was quantified using PUNCH 2-light. Two complementary analyses were performed within the set of 136 matched genes. First, pre-computed mean PUNCH 2-light scores per canonical UniProt entry were compared across the three isoform groups using pairwise Mann–Whitney U tests with BH correction. Second, per-residue PUNCH 2-light scores were used to annotate each residue as TE-derived or non-TE-derived based on TE insertion coordinates projected onto the protein sequence; TE-derived residues of Primate-specific and Pre-Primate isoforms were then compared against all residues of TE-free isoforms from the same genes using pairwise Mann–Whitney U tests with BH correction.

Splice-site strength for all internal exons was computed with MaxEntScan ^105^, producing a score for each 5^′^ donor (5ss) and 3^′^ acceptor (3ss) site. Exons were classified into four categories: TE-derived exons from Primate insertions, TE-derived exons from Pre-Primate insertions, non-TE exons from TE-containing transcripts, and non-TE exons from TE-free transcripts. Two levels of progressive filtering were applied: (1) Considering all isoforms from the 136 matched genes.

For each internal exon, the HEXplorer score (HZEI) was calculated using the precomputed hexamer Z-score table from Erkelenz et al. ^106^. For each 6-nucleotide sliding window across the exon sequence, the HZEI contribution was defined as Z_EI + (−Z_WS), where Z_EI reflects splicing enhancer potential and −Z_WS reflects splicing silencer potential. Per-exon HZEI density was computed as the mean HZEI contribution across all hexamers present in the lookup table, normalizing for exon length. The same group classification and progressive filtering defined for splice-site scoring were applied. Statistical comparisons for both splice-site scores and HZEI density were performed by pairwise Mann–Whitney U tests with BH correction.

## Extended Data Figures

**Extended Data Figure 1.**
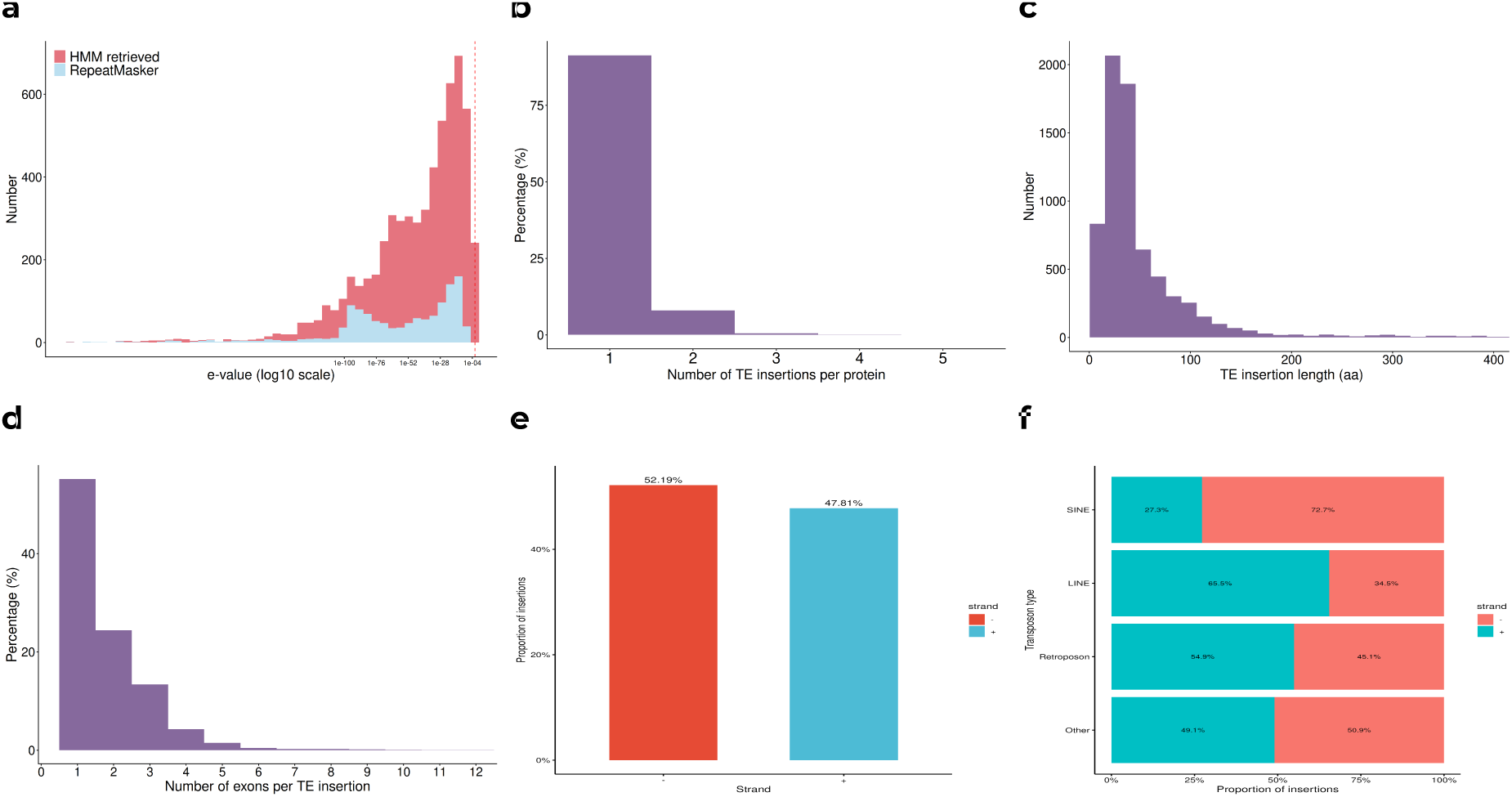
E-value distributions and strand insertion preference. **a.** Comparison of distributions of E-values for each transposon search method. **b,** Proportion of TE insertions per protein. The vast majority of the affected proteins harbor a single insertion event. **c,** TE insertion length distribution at the protein level. **d,** Number of exons involved per insertion event. **e,** Non-redundant transposon insertion hits by strand. Positive (+, n = 3365) and negative (−, n = 3674). **f,** Insertions are shown for each transposon type by strand: SINE (+ n = 315, − n = 840), LINE (+ n = 629, − n = 332), Retroposon (+ n = 50, − n = 41), and Others (+ n = 2,371, − n = 2461).

**Extended Data Figure 2.**
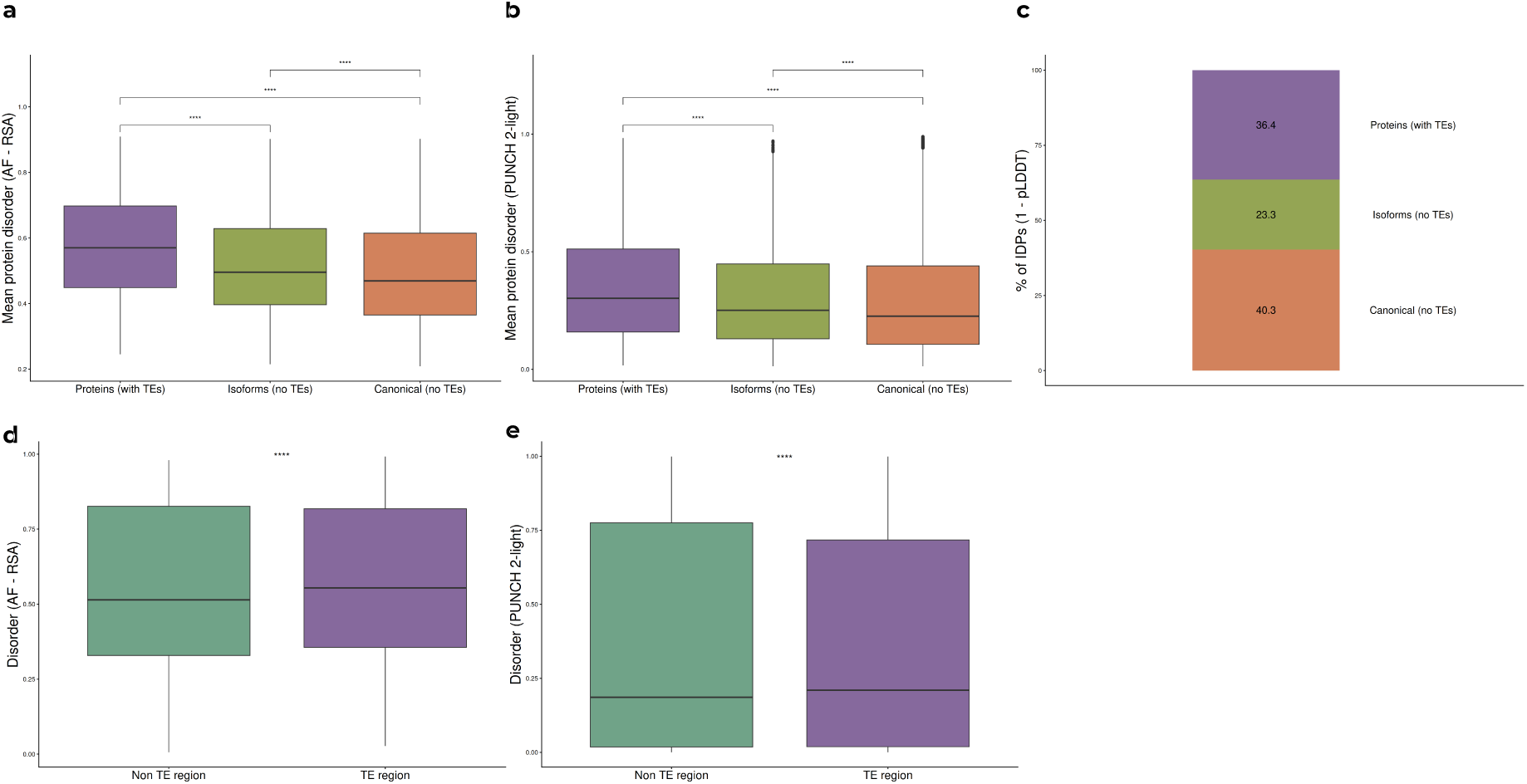
Disorder enrichment in TE-containing proteins is robust across multiple predictors: **a.** Mean protein disorder measured by AlphaFold-RSA across proteins with TE insertions, isoforms without TEs, and canonical proteins without TEs (Mann–Whitney U test, p *<* 0.0001). **b.** Same comparison using PUNCH 2-light disorder scores. **c.** Proportion of IDPs (mean 1 − pLDDT ≥ 0.5) per protein group. **d.** Position-specific disorder measured by AlphaFold-RSA comparing TE-derived and non-TE regions within proteins harbouring TE insertions (Mann–Whitney U test, p *<* 0.0001). **e.** Same comparison using PUNCH 2-light scores. In all panels, significance was assessed by Mann–Whitney U test with Benjamini–Hochberg correction; boxplots show median and interquartile range.

**Extended Data Figure 3.**
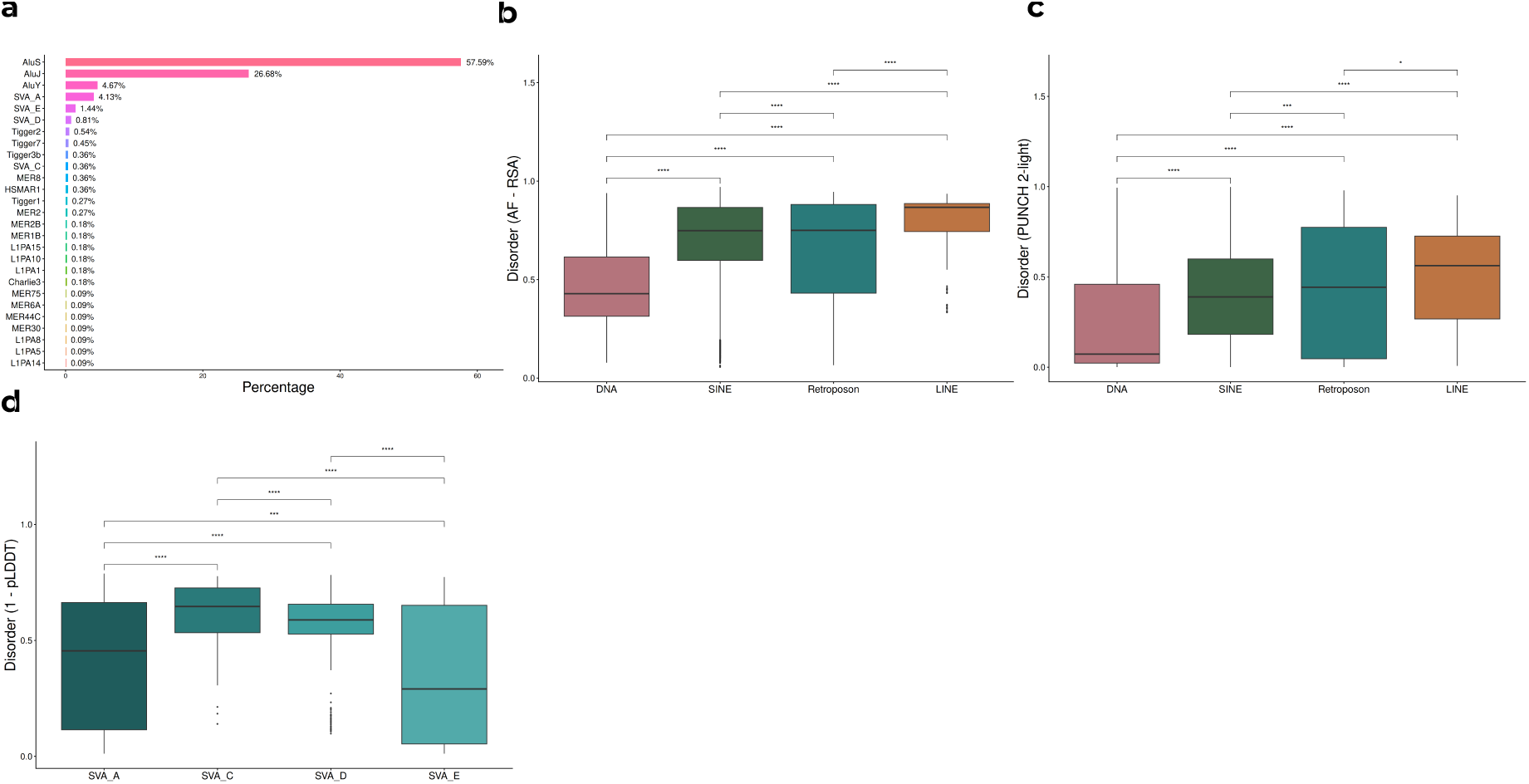
Primate TE insertions show distinct disorder profiles across repeat classes and subtypes: **a.** Distribution of Primate TE insertions by subfamily name, expressed as percentage of total insertions. **b.** Disorder (AF-RSA) across TE classes (DNA, Retroposon, SINE, LINE) for Primate CDS insertion (Mann–Whitney U test, p *<* 0.0001). **c.** Same comparison using PUNCH 2-light disorder scores, confirming the class-specific disorder patterns observed with AF-RSA. **d.** Disorder (1 − pLDDT) across SVA subfamilies (SVA_A, SVA_C, SVA_D, SVA_E) for Primate insertions (Mann–Whitney U test, p *<* 0.0001). In all panels, boxplots show median and interquartile range; significance assessed by Mann–Whitney U test with Benjamini–Hochberg correction.

**Extended Data Figure 4.**
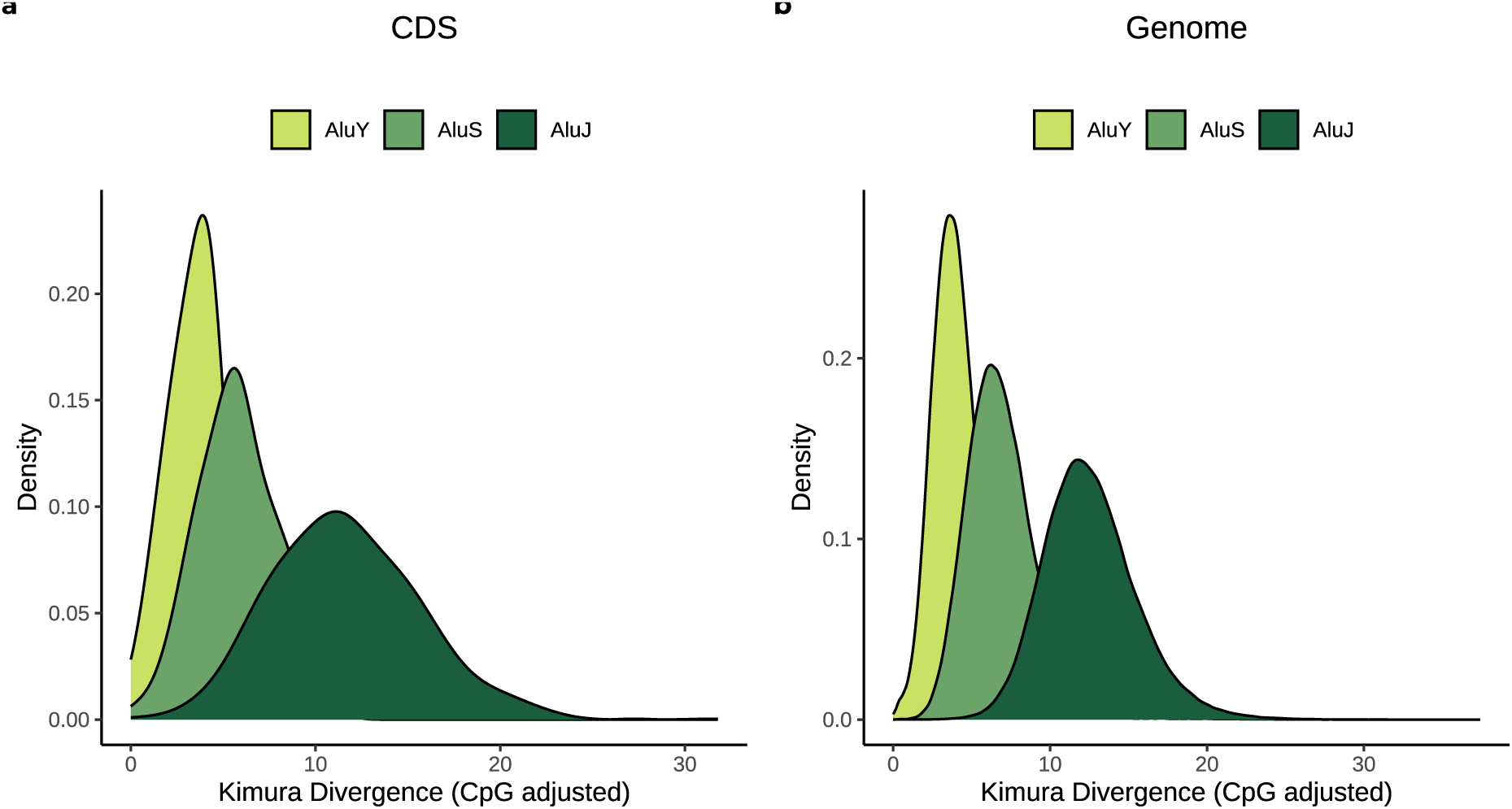
CpG-adjusted Kimura divergence distributions of Alu subfamilies in CDS and genome-wide. Kernel density estimates of CpG-adjusted Kimura divergence for AluY, AluS and AluJ elements in protein-coding sequences (**a**) and across the whole genome (**b**). Numbers of elements per subfamily: **a,** AluY n = 62, AluS n = 724, AluJ n = 353; **b,** AluY n = 133,603, AluS n = 711,041, AluJ n = 297,151. Divergence increases from the youngest (AluY) to the oldest (AluJ) subfamily, consistent with the established evolutionary age ordering.

**Extended Data Figure 5.**
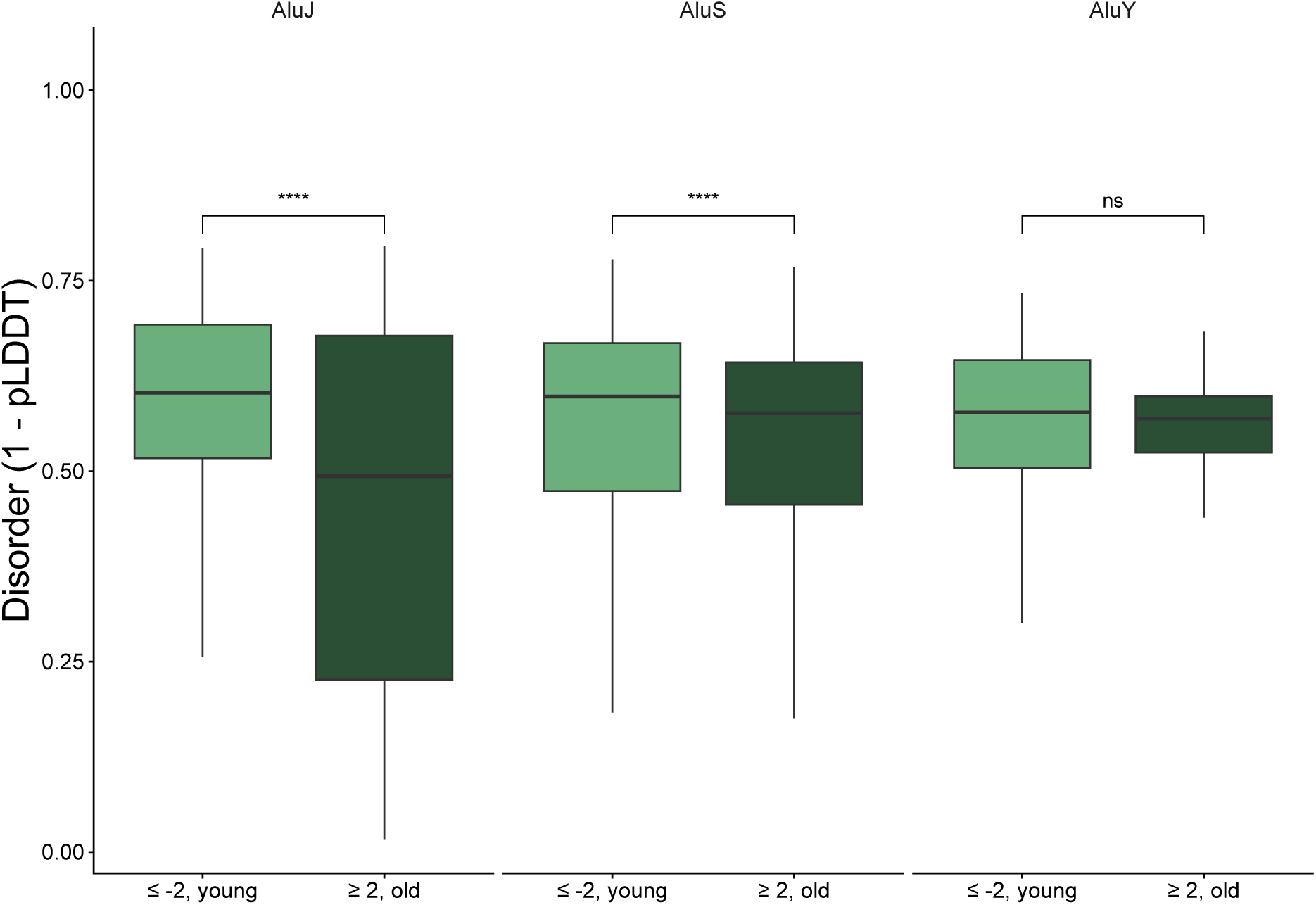
Z-score-based age stratification of Alu subfamily insertions reveals age-dependent disorder differences in AluJ and AluS but not AluY: **a.** Disorder (1 − pLDDT) of CDS-inserted Alu elements classified as younger (Z-score ≤ −2, light green) or older (Z-score ≥ +2, dark green) relative to the genome-wide Kimura CpG divergence distribution of each subfamily. AluJ and AluS Mann–Whitney U test, p *<* 0.0001 and p *<* 0.001, respectively, whereas AluY shows no significant difference (ns). Boxplots show median and interquartile range; significance assessed by Mann–Whitney U test with Benjamini–Hochberg correction. **** p *<* 0.0001; *** p *<* 0.001; ns, not significant.

**Extended Data Figure 6.**
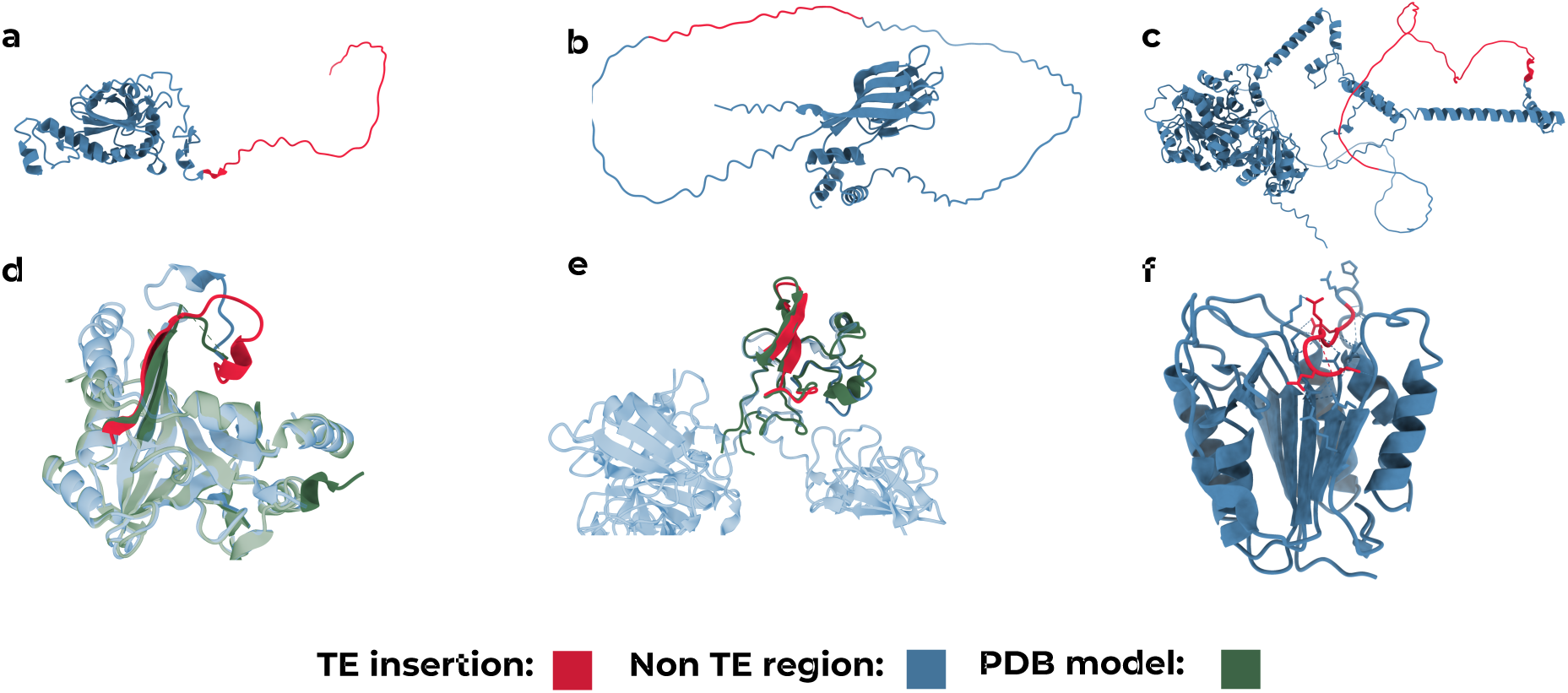
Transposable element insertions contribute to disordered and functionally relevant protein regions: **a.** AlphaFold2 structure of Peroxiredoxin-4 (Q13162) showing an LTR retrotransposon insertion (red, residues 1–45). **b.** AlphaFold2 structure of Nucleophosmin (P06748) with a Helitron DNA transposon insertion (red, residues 159–181). **c.** AlphaFold2 structure of the RNA helicase hPrp28 (Q9BUQ8) showing a LINE element insertion (red, residues 70–140). **d.** Superimposition of Glyoxalase domain-containing protein 4 (Q9HC38; PDB: 3ZI1, green) showing an AluS insertion (red, residues 30–52). **e.** Superimposition of conformational states of tissue-type plasminogen activator (P00750; PDB: 1PK2, green) showing an AluJ insertion (red, residues 269–289). **f.** Structure of MALT1 (Q9UDY8) showing an L1 element insertion (red, residues 493–501) within the elbow-L3 region of the protease active site. In all panels, TE-derived insertions are highlighted in red; non-TE regions are shown in blue (AlphaFold2 models) or green (PDB structures). All visualizations were created with ChimeraX 1.10 (see Methods).

**Extended Data Figure 7.**
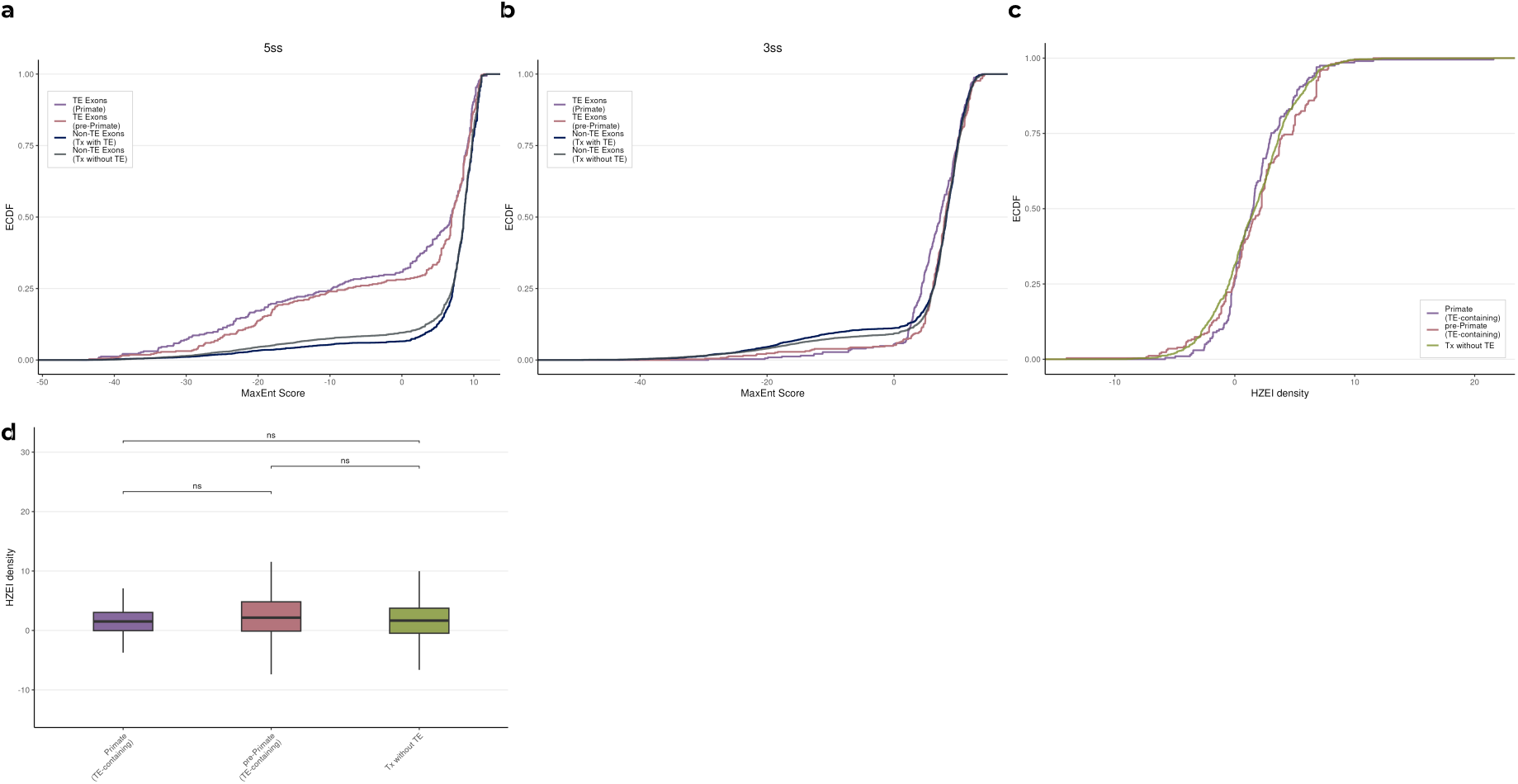
Splicing properties of TE-containing isoforms. **a–b,** Splice-site strength of internal exons in the 136-gene matched set. ECDFs MaxEnt scores of 5^′^ splice-site (5ss, **d**) and 3^′^ splice-site (3ss, **e**). **f–g,** HEXplorer score (HZEI) density of internal exons for Primate-Specific, pre-Primate and transcripts without transposable elements (Tx without TE).

## Notes

https://doi.org/10.5281/zenodo.20648235

### Competing Interest Statement

The authors have declared no competing interest.

